# HIV-1 Vpr drives epigenetic remodeling to enhance virus transcription and latency reactivation

**DOI:** 10.1101/2025.01.31.635859

**Authors:** Nicholas Saladino, Emily Leavitt, Hoi Tong Wong, Jae-Hoon Ji, Diako Ebrahimi, Daniel J Salamango

## Abstract

Despite decades of research, the primary proviral function of the HIV-1 Vpr accessory protein remains enigmatic. Vpr is essential for pathogenesis *in vivo* and for virus replication in myeloid cells, but the underlying cause-and-effect mechanism(s) driving these phenomena are poorly understood. Canonically, Vpr hijacks a cellular ubiquitin ligase complex to target several dozen host proteins for proteasomal degradation. Many of these substrates were recently revealed to be involved in DNA damage repair (DDR), which rationalizes the longstanding observation that Vpr induces constitutive activation of DDR signaling. Here, we use a combination of functional, biochemical, and genetic approaches establish a clear mechanistic link between Vpr-induced DDR signaling and remodeling of the epigenetic landscape to enhance HIV-1 promoter activity during acute infection and virus reactivation from latency. Functional, genetic, and bimolecular fluorescence complementation experiments reveal that Vpr utilizes degradation-dependent and -independent mechanisms to induce epigenetic remodeling and that Vpr segregates into two discrete pools with dedicated activities—A multimeric pool in the nucleus that is associated with chromatin and a monomeric pool associated with DCAF1 in the cytoplasm. Vpr function in remodeling the nuclear environment is present in common HIV-1 subtypes worldwide and provides a mechanistic rationale for its essentiality in virus replication.

**Author summary:** While HIV-1 Vpr plays an essential role in virus replication, the molecular mechanisms underlying its essentiality remain enigmatic. Vpr’s best characterized function is the ability to induce the depletion of several dozen host proteins by hijacking a cellular E3-ubiquitin ligase complex. Here, we establish that Vpr promotes global epigenetic remodeling to enhance HIV-1 promoter activity during acute infection and virus reactivation from latency.

We demonstrate that epigenetic remodeling activity is linked to Vpr’s ability to induce constitutive DNA damage repair signaling, and that it occurs through both degradation-dependent and -independent mechanisms. Moreover, this Vpr function is present in common HIV-1 subtypes circulating globally. This study provides novel mechanistic insights into how HIV-1 exploits host DNA repair pathways and sheds light on Vpr’s proviral function.

## Introduction

HIV-1 encodes four accessory proteins that are essential for virus replication *in vivo*^1–3^. The Vif, Vpu, and Nef accessory proteins promote virus replication by directly counteracting distinct host innate immune defense mechanisms, whereas the primary proviral function of the Vpr accessory protein remains enigmatic^4–6^. Vpr is highly conserved amongst primate lentiviruses and is critical for virus replication in myeloid cells and for pathogenesis *in vivo*^7–11^. Canonically, Vpr engages a host CUL4/DDB1/DCAF1 E3-ubiquitin ligase complex to direct numerous cellular proteins for proteasomal degradation, resulting in systems-level remodeling of the host transcriptome and proteome^12^. Accumulating evidence indicates that a majority of these substrates are involved in DNA repair, DNA modification, or chromatin remodeling^12–14^, which rationalizes observations that Vpr induces constitutive activation of ATM and ATR DNA damage repair (DDR) kinases, the accumulation of DNA strand breaks, and G2/M cell cycle arrest^12,15–18^. Additionally, recent studies revealed that Vif and Vpu independently antagonize diverse DDR pathways to block the activation of antiviral defenses triggered by abnormal DDR signaling or recognition of viral cDNA, signifying that HIV-1 utilizes discrete strategies to fine-tune host DDR responses^19,20^.

Another widely observed Vpr activity is the ability to influence HIV-1 promoter activity in multiple primary and immortalized cell models. For instance, Vpr-deficient viruses exhibit decreased transcription in monocyte-derived macrophages (MDMs) and dendritic cells^9,21–24^. Likewise, Vpr expression leads to enhanced transcription of unintegrated viral DNA and reactivation of virus from latency^25–29^. However, the direct cause-and-effect mechanism(s) underlying Vpr’s ability to modulate virus transcription remains poorly understood. Biochemical and transcriptional studies of HIV-1 infected U2OS and MDMs recently revealed that Vpr-induced DNA damage is correlated with increased NFκB-driven transcription, providing a mechanistic clue for Vpr-directed HIV-1 promoter modulation^18^.

Detection of DNA damage by cellular factors triggers a plethora of dynamic local and global epigenetic post-translational modifications (PTMs) that regulate chromatin accessibility and the binding of DNA repair factors^30–34^. The majority of these modifications involve increased acetylation of lysine residues at the N-terminal tails of core histone proteins. Importantly, several DDR-induced histone PTMs have been associated with increased HIV-1 transcription or reactivation of virus from latency^35^, and multiple lines of evidence have established that decreased acetylation of nucleosomes at the HIV-1 promoter induces latency while increased acetylation is associated with transcriptional activation^36–42^. These observations inspired our hypothesis that Vpr-directed DDR signaling drives epigenetic remodeling to enhance HIV transcription at the integration site. Here, we establish that Vpr induces global epigenetic remodeling in several cell models, including primary MDMs, to enhance HIV-1 promoter activity during acute infection and latency reactivation. Our structure-guided mutagenesis studies demonstrate that Vpr uses a conserved network of electrostatic interactions to induce DDR signaling and epigenetic changes. This comprehensive mutational analysis in combination with chemical inhibitor experiments point to an inseparable relationship between Vpr-induced DDR signaling and epigenetic remodeling activity. Vpr mutants that fail to engage the E3-ligase or genetic depletion of *DCAF1* establish that Vpr utilizes both degradation-dependent and -independent mechanisms to induce epigenetic remodeling. Furthermore, these data suggest that Vpr exists in two functional pools with discrete activities. Functional studies and bioinformatic analyses both indicate that induction of DDR responses and epigenetic remodeling activities are prevalent among common HIV-1 subtypes circulating globally.

## Results

### Vpr induces global epigenetic remodeling through the DNA damage response

To test our hypothesis that Vpr alters the cellular epigenetic landscape, we infected cells with Vpr-proficient (Vpr_WT_) or -deficient (mCh) HIV_NL4-3_ reporter viruses (CMV-driven *mCherry* in place of *nef*) and profiled roughly two-dozen histone PTMs 48 hours post-infection (**Figures 1A** and **1B**). Remarkably, 14 of 17 histone residues exhibited a significant increase in acetylation, phosphorylation, or methylation in differentiated THP1 cells (macrophage-like) infected with Vpr_WT_ virus compared to control infected cells (**Figures 1C**, **1D**, and **S1A**). Infection of primary MDMs and HeLa cells recapitulated these observations, indicating this is not a cell-type specific phenomenon (**Figures 1E**, **S1B**, and **S2**). Importantly, immunoblotting and flow cytometry experiments confirmed Vpr-induced epigenetic changes in differentiated THP1 and HeLa cells following infection (**Figures 1F**, **1G**, **S2B**, and **S2C**). Lastly, time course analyses established that Vpr-directed epigenetic changes persist for ≥96 hours post-infection (**Figure 1H**).

**Figure 1.**
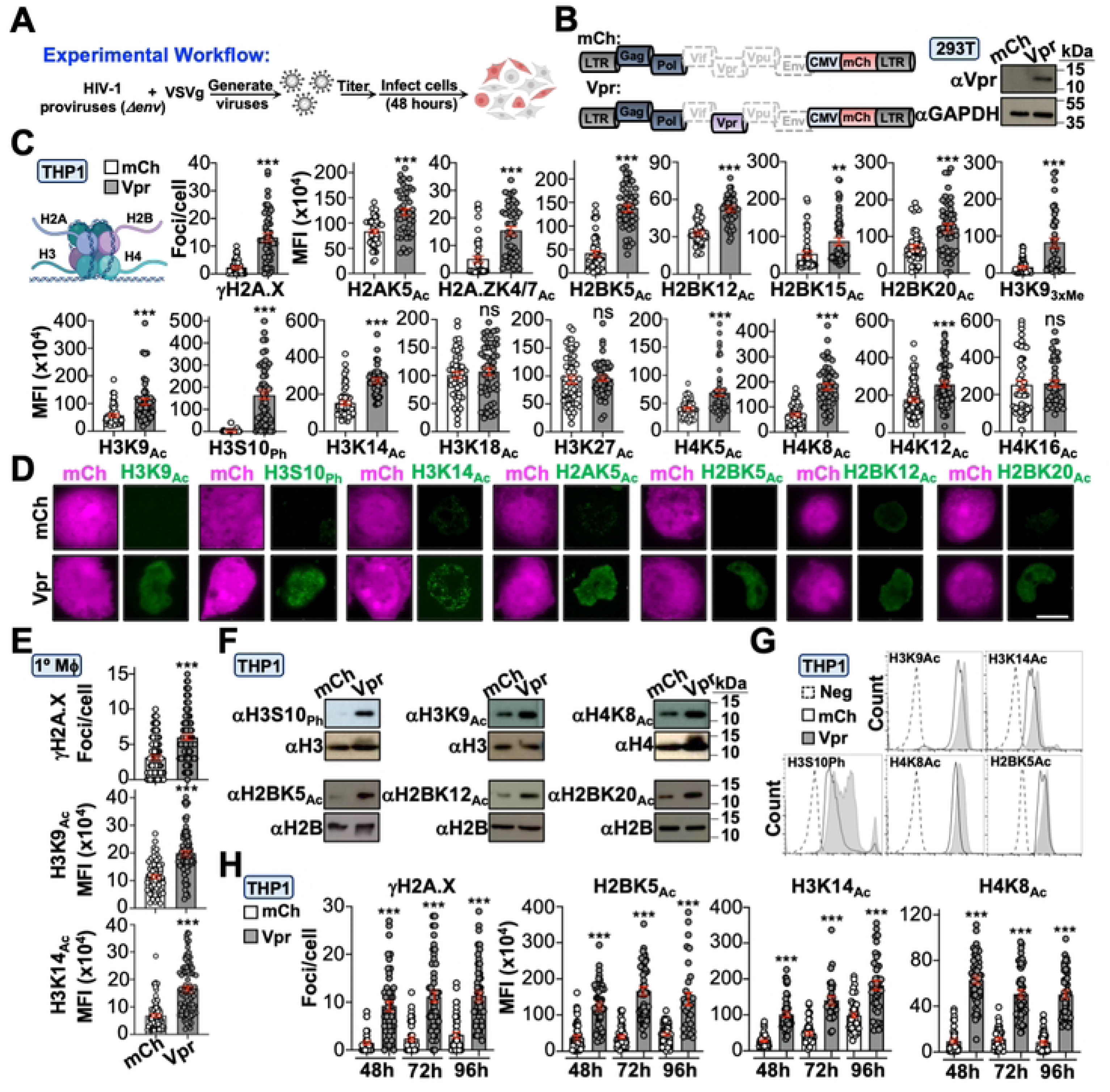
Vpr induces global epigenetic remodeling. (**A** and **B**) Schematic of experimental workflow and validation of Vpr expression in HEK293T cells. For all experiments, cells were infected for 48 hours prior to analysis unless noted. (**C** and **D**) Quantification and representative fluorescence microscopy images of histone marks in THP1 cells infected with indicated viruses (n = 50-75). For all analyses: ns, not significant; ***, p < 0.001; **, p < 0.01; *, p < 0.05; and scale bar = 10 μM. (E) Quantification of DDR activation and histone marks in primary MDMs infected with indicated viruses (n = 75). (F) Immunoblot analysis of histone marks in THP1 cells infected with indicated viruses. (G) Intracellular immunolabeling and flow cytometric analysis of THP1 cells infected with indicated viruses. (H) Time-course analysis of histone marks in THP1 cells infected with indicated viruses (n = 50).

Next, we sought to delineate the upstream mechanism(s) underlying Vpr-directed epigenetic remodeling. As described above, a longstanding observation is that Vpr induces constitutive DDR signaling in multiple cell types (*ex.*, **Figures 1C**, **1E**, **S2D**, and **S2E**). Given that activation of DNA repair is known to alter the epigenetic landscape, we postulated that Vpr’s ability to induce DDR signaling is a prerequisite for epigenetic remodeling activity. To test this, we treated infected cells with caffeine which simultaneously inhibits ATM/ATR and assessed Vpr-induced changes to histone marks. As anticipated, caffeine treatment ablated Vpr’s ability to induce phosphorylation of the DDR marker γH2A.X, and importantly, also ablated the increased abundance of all histone PTMs examined in immunofluorescence microscopy and immunoblotting experiments (**Figures 2A-C**). Moreover, treatment of Vpr_WT_ infected cells with ATM or ATR specific inhibitors recapitulated these observations (**Figure 2B**).

**Figure 2.**
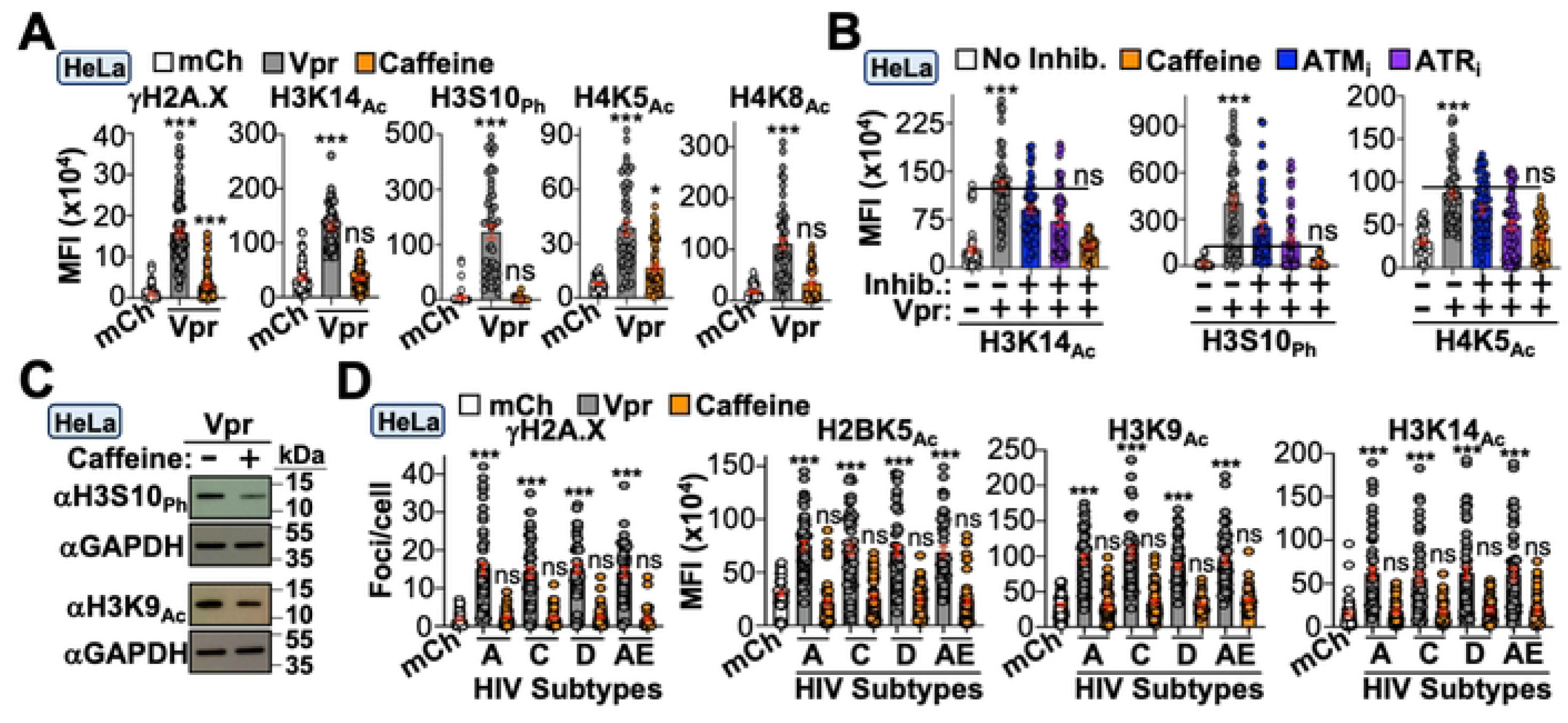
Vpr-induced epigenetic remodeling is dependent on DDR signaling. (A) Quantification of DDR activation and histone marks in HeLa cells infected with indicated virus. Vpr infected cells were treated with vehicle or caffeine (n = 50). (B) Quantification of histone marks in HeLa cells infected with indicated virus in the presence or absence of vehicle, caffeine, ATM_i_, or ATR_i_ (n = 50). (C) Immunoblot analysis of indicated histone marks of Vpr_WT_ infected HeLa cells in the presence and absence of caffeine. (D) Quantification of histone marks in HeLa cells infected with mCh or virus expressing the indicated subtype consensus sequence (n = 50).

Because HIV-1 exhibits a high degree of genetic diversity, we assessed the breadth of this activity across major HIV subtypes circulating globally. Consensus Vpr sequences from subtypes A, C, D, and AE were cloned into a modified version of the HIV_NL4-3_ provirus depicted in **Figure 1A**. To avoid disrupting the *tat* open reading frame, we expressed consensus *vpr* subtypes from a CMV-driven *mCherry-T2A* expression cassette in place of *nef*, which permits independent mCherry and Vpr expression from the same mRNA^43^. HeLa cells were infected with the indicated Vpr subtype and assayed for DDR activation and changes to histone marks. All subtypes were capable of inducing γH2A.X focus formation and increasing the abundance of multiple histone PTMs, which were ablated by caffeine treatment (**Figure 2D**).

### Defining the Vpr surface required for DDR activation and epigenetic remodeling

The co-crystal structure of a Vpr/DCAF1/DDB1 ternary complex revealed that solvent exposed Vpr surfaces exhibit a unique charge dichotomy with electronegative and electropositive N- and C-termini, respectively^44^ (**Figure 3A**, PDB: 5JK7). Because of this, we wondered if the Vpr surface used for inducing DDR signaling and epigenetic remodeling would be electrostatic in nature. We utilized structure-guided mutagenesis to generate a large panel of charge-exchange single amino acid substitution mutations at Vpr surface residues separable from *vif* and *tat* open reading frames. Differentiated THP1 and HeLa cells were infected for 48 hours prior to immunofluorescence microscopy experiments to assess H2A.X and CHK1 phosphorylation (*i.e.,* activation of ATM and ATR signaling pathways, **Figures 3B** and **S3A**). This approach identified two regions of interest: the first encompassed N-terminal electronegative-to-electropositive substitutions that exhibited increased activation of DDR signaling, while the second region involved C-terminal electropositive-to-electronegative substitutions deficient for DDR activation (**Figures 3B** and **S3A**). Importantly, hyperactive (Y15R) and loss-of-function (S79E) mutants exhibited an inextricable link between activation of DDR signaling and modulating histone PTMs in both immunofluorescence microscopy and immunoblotting experiments, further indicating that Vpr-induced DDR signaling precedes epigenetic remodeling activity (**Figures 3C** and **3D**).

**Figure 3.**
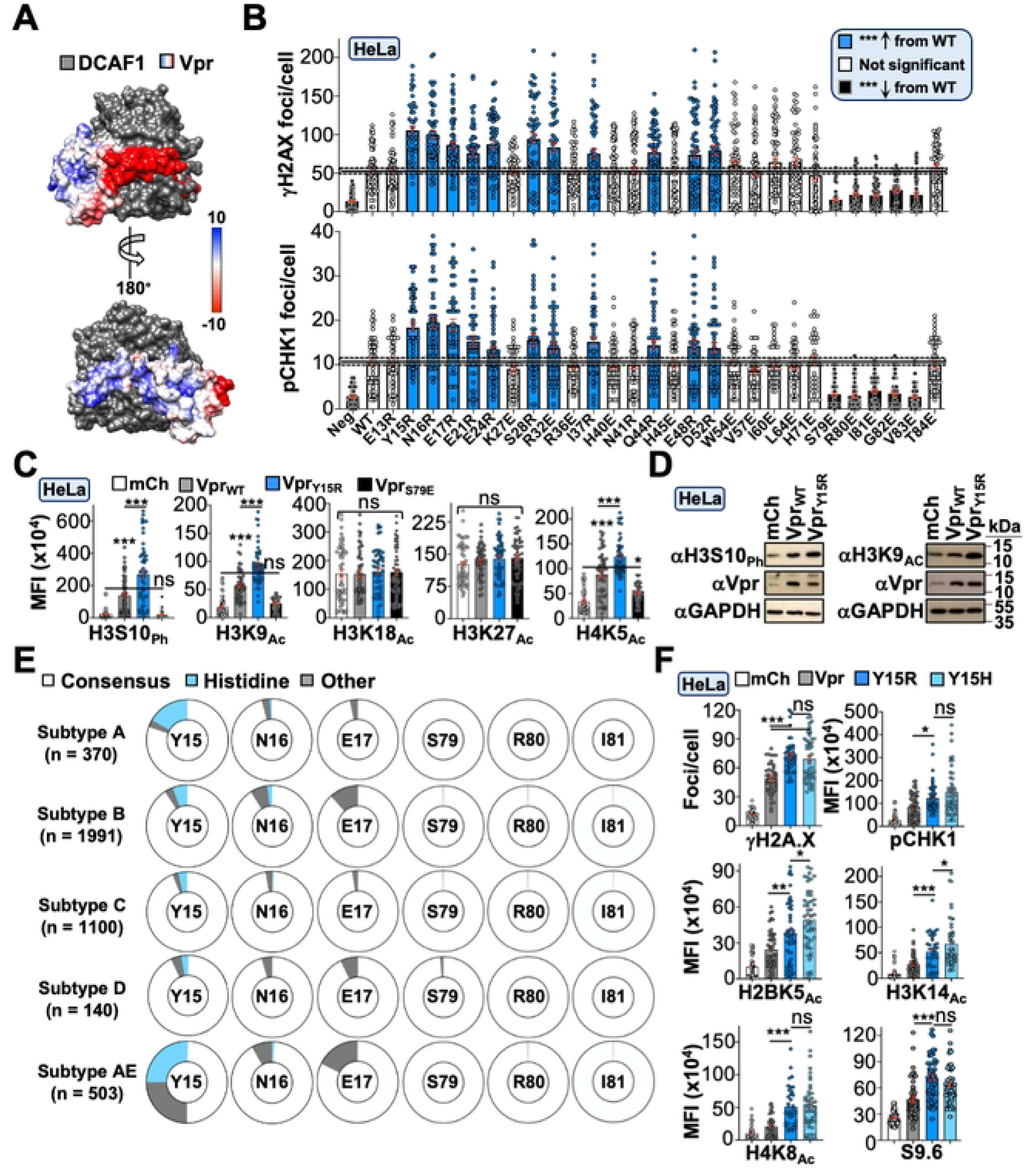
Vpr surface residues used to induce DDR and histone marks. (A) Electrostatic surface potential of a Vpr-DCAF1 co-complex. (B) Quantification of DDR activation following infection of HeLa cells with Vpr_WT_ or mutant viruses (n = 50). (C) Quantification of histone marks in HeLa cells infected with indicated viruses (n = 50). (D) Immunoblot analysis of histone marks in HeLa cells infected with indicated viruses. (E) Circle graphs displaying amino acid variance at relevant positions identified in the mutagenesis screen with consensus residues in the middle of the circle. Sequences obtained from Los Alamos database (n ≈ 4,100). (F) Quantification of DDR activation and histone marks following infection of HeLa cells with Vpr_WT_ or Vpr_Y15H_ mutant (n = 50).

We were intrigued by the hyperactive Vpr mutants and were curious if any are prevalent in patient-derived isolates. Using full-length HIV-1 sequences available in the Los Alamos database (>8,000 bp), we analyzed Vpr polymorphisms at key functional residues when corresponding subtype information was available (∼4,100 sequences total). Our analyses indicated that Vpr residues S79, R80, and I81 were nearly ubiquitously conserved across all major Group M HIV-1 subtypes (**Figure 3E**), which supports functional studies in **Figure 2D**. We also assessed diversity at positions Y15, N16, and E17, and interestingly, the most abundant polymorphism among these residues was a histidine substitution at Y15 (**Figure 3E**). Because histidine mimics the positive charge exchange of the arginine substitution, we reasoned that Y15H would phenocopy Y15R. As anticipated, HeLa cells infected with HIV_NL4-3_ carrying the Vpr_Y15H_ mutation exhibited increased DDR signaling and histone PTMs compared to Vpr_WT_ controls (**Figure 3F**).

### Vpr-directed DDR activation and epigenetic remodeling occur through DCAF1-dependent and -independent mechanisms

While the observations above unambiguously linked Vpr-induced DDR signaling with epigenetic remodeling activity, the molecular mechanism(s) driving these responses were still unclear. Because the best characterized Vpr function is the depletion of host proteins through a CUL4/DDB1/DCAF1 E3-ubiquitin ligase complex, we wanted to determine if E3-engagement is required for inducing DDR signaling and histone PTMs^12^. Proviruses encoding Vpr mutants Q65R or H71R, which are deficient for DCAF1 binding^45,46^, were generated and used to infect HeLa cells. Surprisingly, most histone PTMs, including γH2A.X, only exhibited an ∼20-50% reduction following infection with Vpr_Q65R_ or Vpr_H71R_ viruses compared to Vpr_WT_, suggesting that Vpr utilizes both DCAF1-dependent and - independent mechanisms to modulate DDR signaling and histone PTMs (**Figures 4A** and **S3B**). To further confirm these observations, we depleted *DCAF1* mRNA utilizing two previously validated shRNA constructs^13,47^. Transient expression of knockdown constructs in HeLa cells resulted in robust depletion of DCAF1 protein and mRNA 72 hours post-transfection (**Figure S3C**). Sequential transfection and infection experiments were performed to determine the impact of *DCAF1* depletion on cells infected with Vpr_WT_ or control viruses. Knockdown of DCAF1 recapitulated our findings using Vpr_Q65R_ and Vpr_H71R_ viruses, further supporting the model that Vpr utilizes DCAF1-dependent and - independent mechanisms to drive DDR activation and histone PTMs (**Figure 4B**).

**Figure 4.**
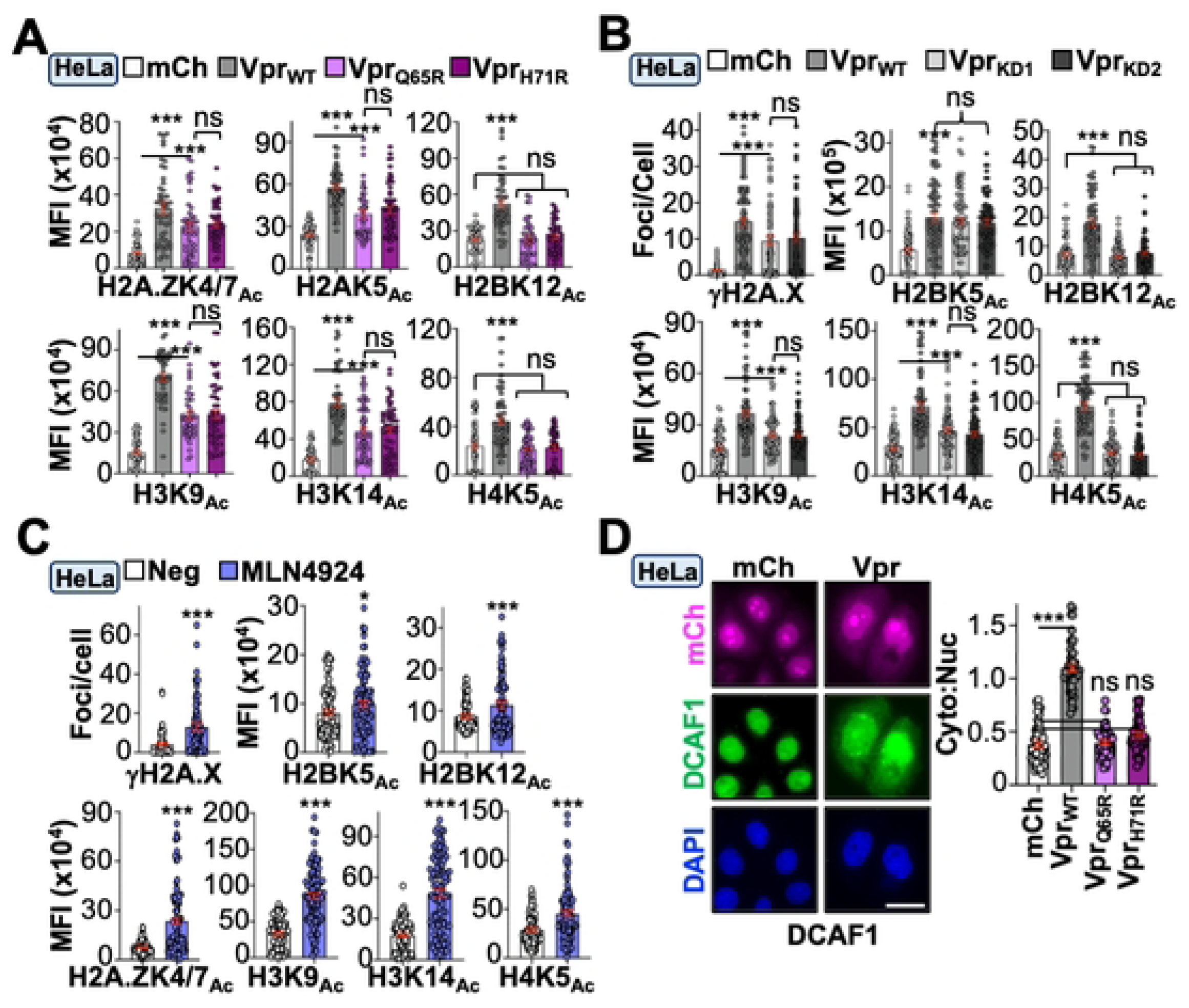
Vpr utilizes DCAF1-dependent and -independent mechanisms to trigger DDR and epigenetic remodeling. (A) Quantification of histone marks in HeLa cells infected with indicated viruses (n = 50). (B) Quantification of histone marks following infection of HeLa cells expressing control or *DCAF1* knockdown constructs (n = 50). (C) Quantification of DDR activation and histone marks in uninfected HeLa cells treated with vehicle or MLN4924 (n = 75). (D) Representative fluorescence microscopy images of DCAF1 localization in HeLa cells infected with indicated viruses and quantification of cytoplasmic:nuclear ratio (n = 50).

A recent study demonstrated that Vpr expression leads to dysregulation of histone 1 ubiquitination and that pharmacological inhibition of the CUL4/DDB1/DCAF1 complex could recapitulate this phenotype in the absence of infection^14^. This raised the possibility that Vpr hijacking the CUL4/DDB1/DCAF1 complex itself may impact the epigenetic landscape. To test this, we assayed γH2A.X focus formation and histone PTMs following treatment of uninfected HeLa cells with the neddylation inhibitor MLN4924, which blocks activation of the CUL4/DDB1/DCAF1 complex^14^. Surprisingly, all histone marks exhibited a significant increase in abundance following MLN4924 treatment, suggesting that E3-engagement by Vpr may also contribute to epigenetic remodeling beyond canonical substrate depletion (**Figure 4C**). Consistent with this possibility, DCAF1 was relocalized from the nucleus to the cytoplasm in HeLa cells infected with Vpr_WT_ virus but not in cells infected with control or Vpr_Q65R_/Vpr_H71R_ viruses (**Figure 4D**). Taken together, these observations support recent studies that suggest Vpr-induced DDR signaling is only partially dependent on DCAF1 engagement^17,48^, and indicate that Vpr utilizes multiple discrete mechanisms to alter the epigenetic landscape.

We next probed DCAF1-independent mechanisms by leveraging the Vpr loss-of-function mutants characterized in **Figure 3**. N-terminally eGFP-tagged wild-type, S79E, R80E, I81E, and G82E Vpr proteins were generated and evaluated using live cell fluorescence microscopy to assess protein abundance and subcellular localization. As depicted in **Figure 5A**, all constructs exhibited whole cell distribution with enrichment at the nuclear envelope, which is consistent with prior studies^49,50^ (**Figure 5A**, left). However, permeabilization of live cells resulted in a complete loss of eGFP fluorescence in cells expressing Vpr mutants but not in cells expressing Vpr_WT_, which exhibited eGFP accumulation at the nuclear envelope and in the nucleoplasm (**Figure 5A**, right and **Figure 5B**). Importantly, loss of eGFP fluorescence in cells expressing Vpr mutants was not due to compromised cell integrity as a co-expressed Nup98-mCherry marker exhibited normal localization at the nuclear envelope (**Figure 5A**).

**Figure 5.**
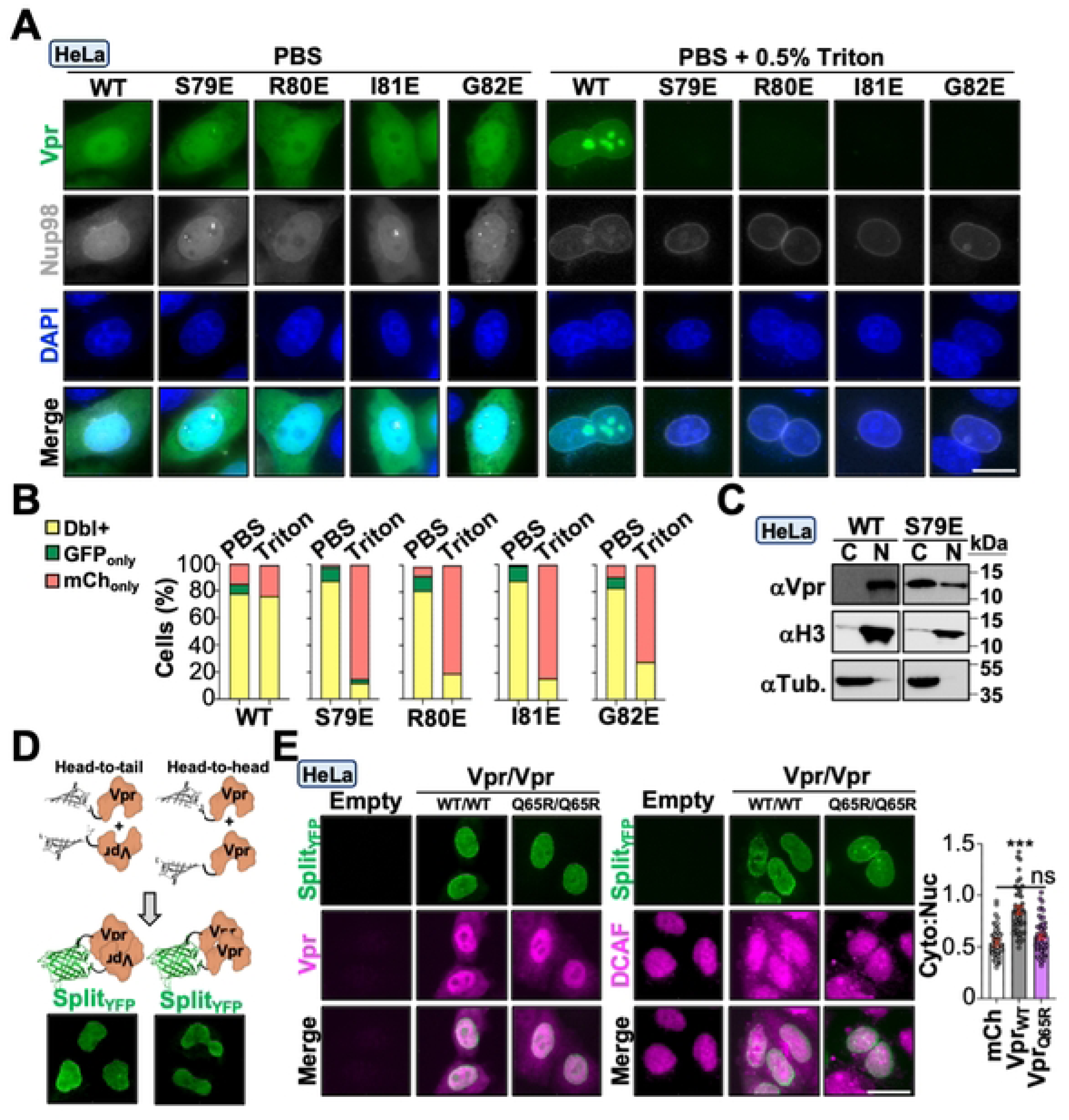
Vpr exists in two functional pools with distinct activities. (A) Representative live cell fluorescence microscopy images of HeLa cells co-expressing indicated eGFP-Vpr constructs and Nup98-mCherry. Cells were either untreated (left) or incubated with 0.5% Triton X-100 (right) prior to imaging. (B) Quantification of dual or singly fluorescence-positive cells (n = 50). (C) Immunoblot analysis of fractionated HeLa cell lysates infected with indicated viruses. (D) Diagrams of BiFC co-transfections and representative live cell fluorescence microscopy images of HeLa cells co-expressing indicated Vpr constructs. (E) Representative fluorescence microscopy images of Vpr (left) and DCAF1 (right) localization in HeLa cells infected with indicated viruses with quantification of DCAF1 cytoplasmic:nuclear ratio (n = 50).

Since surface mapping studies indicated that Vpr may utilize electrostatic interactions to trigger DDR signaling and histone PTMs, we wondered if nuclear retention of Vpr_WT_ was due to chromatin binding, which has been previously observed for recombinant Vpr *in vitro*^51–53^. To test this, we performed chromatin fractionations on HeLa cell lysates infected with Vpr_WT_ or Vpr_S79E_ viruses 48-hours post-infection. As anticipated, Vpr_WT_ was predominantly observed in the nuclear fraction whereas Vpr_S79E_ was enriched in the cytoplasmic fraction, suggesting that nucleus localized Vpr may be binding chromatin (**Figure 5C**). In addition, it has been previously suggested that Vpr’s ability to bind nucleic acids *in vitro* requires oligomerization^51^. We utilized bimolecular fluorescence complementation to determine if nucleus-localized Vpr forms oligomers *in cellulo*. Co-expression of either N- or C-terminally-tagged Vpr proteins fused to split-Venus fragments exhibited robust fluorescence reconstitution at the nuclear envelope and in the nucleoplasm, but not in the cytoplasm (**Figure 5D**).

These observations raised the possibility that Vpr exists in two functionally discrete pools, one associated with chromatin in the nucleus and the other bound to DCAF1 in the cytoplasm. To test this possibility, we leveraged the split-fluorescence system as a live-cell indicator of Vpr localization and function. First, we evaluated Vpr localization in cells that exhibit fluorescence reconstitution (**Figure 5E**). Using a polyclonal antibody that recognizes the Vpr split-Venus fusion protein, we immunolabeled Hela cells transiently expressing Vpr_WT_ and Vpr_Q65R_ proteins; and as anticipated, Vpr was detected in both the nucleus and cytoplasm whereas fluorescence reconstitution only occurred in the nucleus (**Figure 5E**, left). Next, we verified that the Vpr split-Venus fusion protein was capable of relocalizing DCAF1 to the cytoplasm. We observed robust redistribution of DCAF1 to the cytoplasm in cells expressing Vpr_WT_ but not Vpr_Q65R_. As above, cells that exhibited robust DCAF1 relocalization also exhibited robust fluorescence reconstitution (**Figure 5E**, right).

### Vpr induces transcription-coupled R-loops and HIV promoter activity through DDR signaling and epigenetic remodeling

A strong correlation has been established between activation of the DDR response and R-loop accumulation at sites of DNA damage^54–56^. R-loops occur when nascent RNA re-anneals to the transcribed DNA strand, creating a 3-stranded structure containing an RNA/DNA hybrid and an unpaired non-transcribed DNA strand. R-loops generally accumulate at highly transcribed DNA regions and can impede replication forks, potentially leading to DNA strand breaks^57^. We utilized a monoclonal antibody that binds RNA/DNA hybrids with high affinity to determine if Vpr expression correlates with changes in R-loop abundance^58^. HeLa cells and primary MDMs infected with Vpr_WT_ or Vpr_Y15R_/Vpr_Y15H_ viruses exhibited a significant increase in R-loop abundance compared to control infected cells 48-hours post-infection (representative images in **Figure 6A**, quantification in **Figures 6B** and **3F**). To confirm these were *bona-fide* R-loops, infected cells were treated with RNase H which specifically degrades the RNA component of RNA/DNA hybrids^58,59^. Both control and Vpr_WT_ infected cells exhibited a significant reduction in immunostaining compared to vehicle treated controls, indicating these were authentic R-loops (**Figure 6C**). We also tested if R-loops were associated with transcription by treating infected cells with the transcriptional inhibitor triptolide^60^. As depicted in **Figure 6D**, triptolide treatment significantly reduced R-loop abundance in both control and Vpr_WT_ infected cells, suggesting these were occurring co-transcriptionally. We also investigated the requirement for DCAF1-engagment and DDR signaling/epigenetic remodeling activity in R-loop accumulation. Vpr_Q65R_-infected cells exhibited an ∼50% reduction in R-loop abundance whereas cells infected with Vpr_S79E_ virus exhibited levels comparable to control infected cells, indicating that R-loop formation occurs through both DCAF1-dependent and -independent mechanisms (**Figure 6E**). Furthermore, inhibition of DDR signaling through either caffeine treatment or ATM/ATR-specific inhibitors also ablated Vpr-induced R-loops, suggesting these phenotypes are tightly associated (**Figure 6F**).

**Figure 6.**
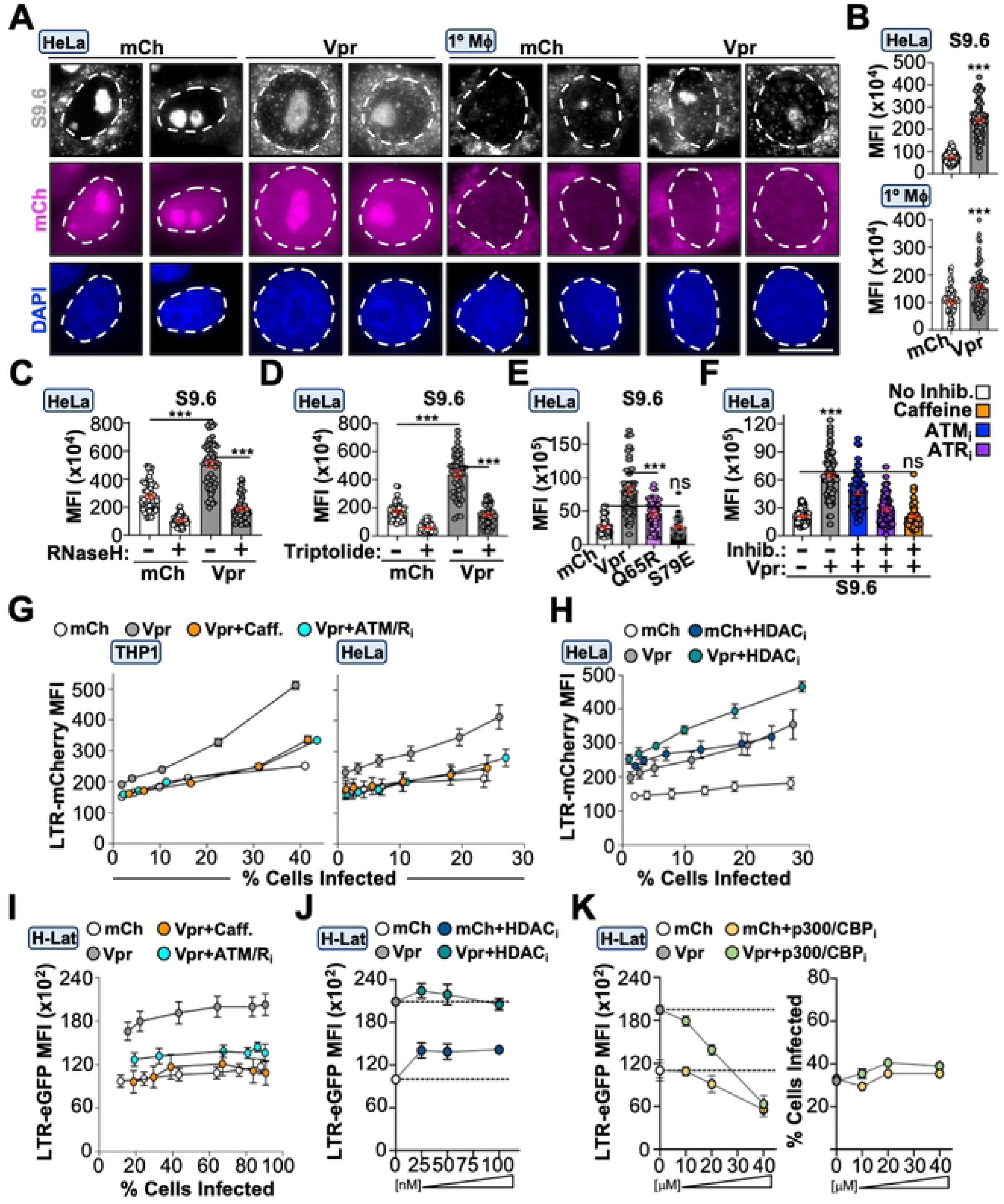
Vpr induces transcription-coupled R-loops and HIV promoter activity through the DDR response. (**A** and **B**) Representative fluorescence microscopy images (left) and quantification (right) of R-loop abundance in Hela and primary MDMs infected with indicated viruses (n = 50). (**C-F**) Quantification of R-loop formation in HeLa cells infected with indicated viruses. Samples were treated with RNaseH (C), triptolide (D), untreated (E), or treated with DDR inhibitors (F), respectively (n = 50). (G) Flow cytometric analysis of HIV LTR activity in HeLa cells (left) and THP1 cells (right) infected with increasing MOI of indicated viruses in the presence or absence of DDR inhibition. (H) Flow cytometric analysis of HIV LTR activity in HeLa cells infected with increasing MOI of indicated viruses in the presence or absence of HDAC inhibition. (**I-K**) Flow cytometric analysis of HIV LTR reactivation following infection with increasing MOI of indicated viruses in the presence or absence of indicated inhibitors.

To determine the virologic function of these Vpr activities, we generated proviruses that express mCherry from the native HIV-1 promoter. Differentiated THP1, HeLa, or primary MDMs infected with Vpr_WT_ virus exhibited significantly increased mCherry expression compared to control infected cells at all doses tested (**Figures 6G**, **S4A**, and **S4B**). Importantly, treatment with caffeine or ATM/ATR-specific inhibitors reduced Vpr-enhanced mCherry expression to that of control infection (**Figures 6G**, **S4A**, and **S4B**). Because recent studies have suggested that Vpr antagonizes class I histone deacetylase enzymes (HDACs), and HDAC-inhibitors (HDAC_i_) are well-known latency-reactivating agents (LRAs)^27,41,42,61–63^, we assessed HIV-1 promoter activity in the presence and absence of HDAC inhibition. As expected, control infected cells treated with HDAC_i_ exhibited a significant increase in mCherry expression compared to vehicle treated controls (**Figure 6H**). Interestingly, we observed a further increase of mCherry fluorescence in Vpr_WT_-infected cells treated with HDAC_i_ (**Figure 6H**), suggesting that HDAC antagonism is not the primary mechanism through which Vpr modulates the epigenetic landscape during acute infection, even though Vpr-induced histone marks are sensitive to HDAC_i_ treatment (**Figure S4C**).

Accumulating evidence indicates that Vpr can reactivate virus from latency^27–29,64,65^. Since Vpr-induced DDR activation correlated with enhanced HIV-1 promoter activity during acute infection, we reasoned that it would also promote latency reversal. To test this, we generated a clonal “off-to-on” HeLa latency model (H-Lat) that stably expresses the HIV 5’-LTR promoter upstream of an eGFP expression cassette, allowing for direct correlation between promoter activation and Vpr expression in the absence of *tat* or *vpr* expression from an integrated provirus. Under steady-state conditions, eGFP expression is not detectable by fluorescence microscopy or flow cytometry (**Figure S4D** and **S4E**). H-Lat cells were infected with increasing amounts of Vpr_WT_ or control viruses and eGFP expression was assessed via fluorescence microscopy and flow cytometry (**Figures 6I**, **S4D**, and **S4E**). Infection of H-Lats with Vpr_WT_ virus induced significantly higher eGFP expression compared to control infections, regardless of MOI, which could be ablated by caffeine or ATM/ATR-specific inhibitors (**Figure 6I**). Interestingly, treating H-Lats with HDAC_i_ resulted in a significant increase in eGFP expression for control infected cells but not for Vpr_WT_ infected cells (**Figure 6J**). We reasoned that Vpr could be saturating activation of the integrated 5’-LTR reporter, or, in this context Vpr is primarily inducing expression through histone acetylation. Regardless, HDAC_i_ treatment of control infected cells did not induce eGFP expression to the level of Vpr_WT_ infected cells, suggesting that an additional mechanism is involved. Because the histone acetyltransferase p300/CBP has been implicated in enhancing HIV promoter activity^37,38,66,67^, we treated infected cells with increasing concentrations of a p300/CBP inhibitor. Cells infected with Vpr_WT_ were more sensitive to inhibitor treatment compared to control infected cells, suggesting that p300/CBP may also be involved in Vpr-enhanced promoter activation (**Figure 6K**, left). Importantly, decreased eGFP expression was not due to cytotoxicity as infection rates and cell survival were unchanged at the highest p300/CBP inhibitor concentration (**Figure 6K**, right).

## Discussion

While Vpr has been studied extensively, direct cause-and-effect mechanism(s) underlying its proviral function are unclear. Here, we demonstrate that Vpr’s ability to induce DDR signaling promotes epigenetic remodeling and enhances HIV-1 promoter activity during acute infection and latency reactivation. Vpr mutants that are hyperactive or deficient for DDR activation exhibit analogous phenotypes for epigenetic remodeling and modulating HIV-1 promoter activity, suggesting a direct cause-and-effect relationship. Vpr determinants required for these activities were mapped to distinct surface regions that are electrostatic in nature and likely promote chromatin association to induce remodeling of the nuclear environment. Biochemical, genetic, and pharmacologic experiments indicate that Vpr utilizes both degradation-dependent and - independent mechanisms to induce epigenetic remodeling and likely exists as two discrete functional pools. Functional studies and phylogenetic analyses combine to indicate mechanistic conservation in common HIV-1 subtypes circulating globally.

Here, our findings support previous observations that modulating histone acetylation directly influences HIV-1 promoter activity. We demonstrate that Vpr induces DDR-dependent acetylation of several histone residues known to be dynamically modulated during DDR^30–34,68^ and are associated with transcriptionally active euchromatin^30,69^. Moreover, recent studies indicate that Vpr induces the depletion and chromatin-disassociation of class I HDACs^27,64^ and can directly engage the p300/CBP acetyltransferase^70–72^. These observations are consistent with reports demonstrating that p300/CBP and GCN5 promote HIV-1 transcription^38,66,67^, and that pan-HDAC inhibitors reactivate latent reservoirs^41,42,61–63^. Furthermore, functional studies revealed that Vpr expression and HDAC_i_ treatment synergize to further enhance HIV-1 promoter activity. Because current LRAs are inefficient at reactivating diverse viral reservoirs *in vivo*, future studies that carefully define epigenetic synergy between Vpr and HDAC inhibitors may inform the development of novel LRAs.

While Vpr is thought to primarily function through the depletion of host factors, emerging evidence supports a model wherein Vpr also functions independent of E3-ligase engagement. For instance, structural studies have demonstrated that Vpr binds the nucleotide-excision repair protein RAD23A using the same interface as DCAF1 and that binding is mutually exclusive^73^. Additionally, Vpr mutants that fail to bind DCAF1 or deplete canonical DNA repair substrates can still induce DNA strand breaks, activate DDR signaling, and exert a proviral effect during acute infection^17,18,74^. These observations raise the possibility that Vpr exists in at least two functional pools that have distinct activities. Our findings further support this model by indicating Vpr-induced epigenetic remodeling occurs through DCAF1-dependent and independent mechanisms, and that nucleus localized Vpr is multimeric while cytoplasmic Vpr likely exists as soluble monomers. Furthermore, fractionation, localization, and live-cell permeabilization experiments combine to suggest that cytoplasmic Vpr is DCAF1 bound and can diffuse freely while nucleus-localized Vpr is likely associated with nucleic acid.

The *vpr* gene is conserved in all primate lentiviral lineages and is required for pathogenesis in humans and primates^7,11^. In limited testing, Vpr isolates from some HIV-2 and SIV strains have been shown to activate DDR signaling and induce DNA strand breaks, suggesting that epigenetic remodeling activity may be evolutionarily conserved^17,75^. These observations rationalize previous findings that caffeine treatment can inhibit HIV-1 replication *in vitro* and *in vivo*^76–79^. Furthermore, functional and bioinformatic analyses of common HIV-1 Group M subtypes that indicate epigenetic remodeling activity is prevalent. Taken together, these observations support the notion that these Vpr activities are advantageous for viral pathogenesis.

## Materials and Methods

### Experimental replicates, quantification, and statistical analyses

All experimental procedures were repeated at least three independent times except when using primary MDMs which were performed using PBMCs from two independent donors. All flow cytometry data were analyzed using FlowJo v10 software. Focus formation and mean fluorescence intensity (MFI) analyses were calculated using ImageJ software and analyzed using GraphPad Prism 6 software. Briefly, boundaries of infected cell nuclei were defined using DAPI staining as an indicator and then foci were quantified using the “find maxima” feature and eGFP mean fluorescence intensity was defined by analyzing integrated pixel intensity of the defined nuclear area minus the background signal intensity of an adjacent area with identical dimensions. The Vpr-DCAF co-structure depicted in **Figure 2** was generated using Chimera protein modeling software (PDB: 5JK7). Statistical analyses were performed using either an unpaired two-tailed Student’s *t-*test or a one-way ANOVA in GraphPad Prism 8 after confirming that all data followed a normal distribution. Schematics were generated using BioRender.

### Plasmids and cloning

Proviral expression plasmids have been described previously^80^. For *DCAF1* knockdown, previously validated shRNA sequences were cloned into a pLKO plasmid expressing mTag-BFP2 in place of puromycin (KD_1_, GCTGAGAATACTCTTCAAGAA; KD_2_, TCACAGAGTATCTTAGAGA)^13,47^. eGFP-tagged Vpr proteins and Nup98-mCherry were cloned into a pcDNA 5TO expression vector. Vpr single amino acid substitution mutants were generated by PCR amplification using Phusion high fidelity DNA polymerase (NEB, Ipswich, MA) and overlapping PCR. All constructs were confirmed by Sanger sequencing and restriction enzyme digestions.

### Cell lines and culture conditions

HeLa and HEK293T cells (American Type Culture Collection) were maintained in DMEM medium (Gibco cat #11-965-118) supplemented with 10% fetal bovine serum (FBS; Gibco, Gaithersburg, MD) and 0.5% penicillin-streptomycin (50 units; Gibco, Gaithersburg, MD). HeLa/HEK293T cells were transfected using 1mg/ml polyethylenimine (PEI; Fisher #NC1014320) at a ratio of 3 µL per 1 µg of DNA. For virus generation, HEK293T cells were co-transfected with a VSV-G expression vector along with the indicated proviral plasmid. Medium was collected 48 h post-transfection and frozen at ^-^80 degrees C until used. THP1 cells were maintained in RPMI medium (Gibco; cat #11-875-093) supplemented with 10% FBS and 0.5% penicillin-streptomycin. For THP1 differentiation, cells were incubated with 50-70 ng/ml of phorbol 12-myristate 13-acetate (PMA; Sigma #P8139) for 48-96 h. Primary MDMs were isolated via plastic adhesion of human peripheral blood mononuclear cells (hPBMCs; Lonza cat #CC-2704). To stimulate differentiation, cells were incubated with 50 ng/µl of granulocyte-macrophage colony-stimulating factor (GM-CSF; PeproTech cat #300-03) for 72-96 h until macrophage morphology was observed. For generating the HIV LTR-GFP cell line, HeLa cells were stably transfected with a previously described 5’-LTR eGFP expression vector for 48 h before selecting with 1mg/ml puromycin (Thermo Fisher, BP295100) to generate pure cell populations^81^. Clonal isolates were generated using limiting dilution in 96-well flat bottom culture plates and validated for reactivation efficiency using HDAC_i_ treatment.

### Fluorescence microscopy and immunostaining

For immunofluorescence microscopy experiments using HeLa cells, approximately 5,000 cells were seeded into a 96 well glass-bottom imaging plate (Ibidi #89627) and allowed to adhere overnight at 37 degrees C. The next day, cells were infected with the indicated virus for 48 h prior to being washed 1x with PBS and then fixed using 4% paraformaldehyde (PFA) for 10 min at room temperature. After fixation, cells were washed 3x with PBS in 5-minute intervals and permeabilized using PBS plus 0.3% Triton X-100 (PBST) for 10 minutes at room temperature. Cells were blocked using PBST supplemented with 5% bovine serum albumin (BSA, Fisher Bioreagents BP9703100), 10% goat serum (Sigma-Aldrich, G9023-10mL) and 0.3 M glycine for 2 hours at room temperature while rocking. After blocking, samples were incubated with primary antibodies against γH2A.X (1:300, Cell Signaling 9718), pCHK1 (1:50, Cell Signaling 2348), pCHK2 (1:200, Cell Signaling 2661), acetyl-histone H2AK5 (1:1000, Cell Signaling 2576), acetyl-histone H2A.Z K4/K7 (1:1000, Cell Signaling 75336), total H2B (1:1000, Cell Signaling 12364), acetyl-histone H2BK5 (1:1000, Cell Signaling 12799), acetyl-histone H2BK12 (1:1000, Cell Signaling 5410), acetyl-histone H2BK15 (1:1000, Cell Signaling 9083), acetyl-histone H2BK20 (1:1000, Cell Signaling 34156), total H3 (1:1000, Cell Signaling 4499), phospho-histone H3S10 (1:1000, Cell Signaling 53348), acetyl-histone H3K9 (1:1000, Cell Signaling 9649), tri-methyl-histone H3K9 (1:1000, Cell Signaling 13969), acetyl-histone H3K14 (1:1000, Cell Signaling 7627), acetyl-histone H3K18 (1:1000, Cell Signaling 13998), acetyl-histone H3K27 (1:1000, Cell Signaling 8173T), tri-methyl histone H3K27 (1:1000, Cell Signaling 9733), total H4 (1:1000, Cell Signaling 2935), acetyl-histone H4K5 (1:1000, Cell Signaling 8647), acetyl-histone H4K8 (1:1000, Cell Signaling 2594), acetyl-histone H4K12 (1:1000, Cell Signaling 13944T), DCAF1/VprBP (1:400, Proteintech 11612-1-AP) or S9.6 DNA-RNA hybrid (1:200, Kerafast ENH001) in blocking buffer overnight at 4 degrees C. The next day, cells were washed 3x with PBS in 5-minute intervals and then incubated with anti-mCherry conjugated to Alexa Fluor 594 (1:800, Invitrogen M11240), secondary anti-rabbit-IgG conjugated to Alexa Fluor 488 (1:800, Cell Signaling 4412), secondary anti-rabbit-IgG conjugated to Alexa Fluor 594 (1:800, Cell Signaling 8889) or secondary anti-mouse-IgG conjugated to Alexa Fluor 488 (1:800, Cell Signaling 4408) in blocking buffer at room temperature for 1 h. After incubation, cells were washed 3x with PBS at 5-minute intervals and stained with NucBlue stain (Thermo Fisher, R37605) and imaged. An EVOS M500 fluorescence microscope was used for imaging, using 60x or 100x oil-immersion objectives. For immunofluorescence microscopy experiments using differentiated THP1 cells, approximately 20,000 cells were seeded into a 96-well glass-bottom imaging plate in the presence of 50-70 ng/ml PMA and co-infected with the indicated virus for 48-96 h prior to being subjected to immunofluorescence microscopy. For immunofluorescence experiments using primary MDMs, approximately 15,000 macrophages were seeded into a 96-well glass bottom imaging plate 5 days post differentiation with GM-CSF. The next day, cells were infected for 48 h prior to being prepared for immunofluorescence microscopy.

### Inhibitor treatments

For inhibitor experiments, cells were treated 24-hours post-infection with either 10 nM ATM inhibitor (Fisher, #AZD1390), 10 µM ATR inhibitor (Fisher, #NU6027), 3 mM caffeine, 10 µM mocetinostat (SelleckChem, cat #S1122), 50 nM panobinostat (Sigma, SML3060), or 10µM C646 (SelleckChem, cat #S7152) for 24 hours. Cells were then processed for downstream analysis as indicated.

### Live cell permeabilization experiments

Approximately 5,000 HeLa cells were seeded into a 96-well glass-bottom imaging plate and allowed to adhere overnight at 37 degrees C. The next day, cells were co-transfected with approximately 150 ng of pcDNA eGFP-Vpr wild-type or mutant constructs and 150 ng of pcDNA Nup98-mCherry. 48 h post-transfection, live cells were treated with 0.5% Triton X-100 in PBS for 5 minutes and imaged using an EVOS M5000 microscope.

### Immunoblotting

For immunoblotting experiments, HeLa or THP1 cells were plated at a seeding density of 350,000 cells per well in a 12-well tissue culture plate and infected with the indicated virus for 48 h prior to harvesting. Cell pellets were resuspended in RIPA buffer (50 mM Tris [pH 8.0], 150 mM NaCl, 1 mM ꞵ-mercaptoethanol, 1% Triton X-100, 0.1% SDS, 0.5% deoxycholate) supplemented with a protease and phosphatase inhibitor cocktail (Thermo Scientific #78440). Cell lysate was combined with 5x loading dye (62.5 mM Tris [pH 6.8], 20% glycerol, 5% ꞵ-mercaptoethanol, 2% SDS, 0.05% bromophenol blue) and samples were separated using a 12% SDS-PAGE gel and transferred using a 0.2 µM polyvinylidene fluoride (PVDF) membrane (Thermo Scientific #78440). Membranes were blocked in 5% BSA in PBST for 1 h and then incubated with primary antibodies described above, and antibodies against Vpr (1:500, Proteintech 51143-1-AP) and tubulin (1:1000, Cell Signaling 3873) diluted in blocking buffer overnight at 4 degrees C while rocking. The next day, primary antibody was removed and membranes were washed 3 x with PBST for 5 minutes each. After a brief 30 minute blocking step with 5% milk in PBST, membranes were incubated with ɑ-mouse HRP (1:10000, SantaCruz sc-525409) or ɑ-rabbit HRP (1:10000, Cell Signaling 7074P2) secondary antibody diluted in 5% milk PBST and rocked for 1 h at room temperature. After washes with PBST, blots were incubated with West Pico PLUS chemiluminescent substrate (Thermo Scientific #34580) for 5 minutes before imaging on a BioRad ChemiDoc™ MP Imaging System.

### Bimolecular fluorescence complementation

Vpr was cloned into a pcDNA expression vector expressing CMV-driven N- or C-terminal fragments of Venus (24947387). This expression vector also encoded SV40-driven mCherry in place of hygromycin as a transfection reporter. Approximately 5,000 HeLa cells were plated into a 96-well glass-bottom plate and allowed to adhere overnight. The next day, cells were transfected with 150 ng each of N- or C-terminally tagged Vpr derivatives and imaged 24-hours post transfection. For coupling immunofluorescence microscopy with bimolecular fluorescence complementation, 48-hours post transfection HeLa cells were fixed with 4% PFA and immunofluorescence labeling was performed as described above (DCAF1, 1:400, Proteintech 11612-1-AP; GFP, 1:500, Cell Signaling 2956).

### Chromatin fractionation

For fractionation experiments, SimpleChIP® Enzymatic Cell Lysis Buffers A and B (Cell Signaling 14282) were utilized. Briefly, cells were infected as described above and trypsinized, washed, and fixed in 1% paraformaldehyde for 30 minutes. After washing in cold PBS, cells were centrifuged and resuspended in cold Buffer A supplemented with 1M dithiothreitol (DTT) and 200x protease inhibitor cocktail (PIC; Cell Signaling 7012) and incubated on ice for 10 minutes, with mixing by inversion every 3 minutes. Extracted intact nuclei were pelleted by cold centrifugation at 2,000 RCF for 5 minutes, the cytoplasmic fraction was removed and transferred to a fresh Eppendorf tube. Intact nuclei were washed with Buffer A and resuspended in ice-cold Buffer B supplemented with 1M DTT, pelleted at 2,000 xg for 5 minutes. This process was repeated twice. To fragment genomic DNA, nuclei were treated with micrococcal nuclease (MNase; Cell Signaling 10011) and incubated for 30 minutes at 37℃ with frequent mixing afterwhich this reaction was stopped using 0.2% sodium dodecyl sulfate (SDS) and 10mM ethylenediaminetetraacetic acid (EDTA) stop buffer for 10 minutes at 37℃. The nuclear fraction was then clarified by cold centrifugation at 9,400 xg for 10 minutes and both the cytoplasmic and nuclear fractions were subjected to immunoblotting.

### Flow cytometry

For immunolabeling to assess histone abundance, the indicated cell lines were seeded into 12-well plates and adhered overnight prior to being infected with the indicated virus. At 48 h post-infection, cells were washed and harvested following trypsinization, or collected in suspension for non-differentiated THP1 cells, and centrifuged at 500 RCF for 10 minutes. After washing with PBS, cells were centrifuged and resuspended in 4% PFA for 30 minutes, washed 3x with PBS, and resuspended in PBST and incubated at room temperature for 30 minutes. Cells were then resuspended in 5% BSA/PBST blocking buffer, incubated for 1 hour, pelleted via centrifugation at 500 RCF, and resuspended in primary antibody directed against the indicated histone mark. Samples were washed 3x times with PBS and resuspended in secondary anti-mCherry Alexa Fluor 594 and secondary anti-rabbit-IgG Alexa Fluor 488 in blocking buffer at room temperature for 1 h. Cells were washed with PBS, resuspended in cold PBS and subjected to flow cytometry using an Invitrogen Attune NxT flow cytometer.

For analysis of HIV-1 LTR activity, HeLa, THP1, or primary MDMs were seeded as described above and infected with the indicated virus for 24 hours. The next day, cells were treated with vehicle or the indicated inhibitor for 24 hours. At 48 h post-infection, cells were washed once with PBS, detached using Trypsin/EDTA or Accutase, and centrifuged at 500 RCF for 10 minutes at 4 degrees. Cells were washed 1x in PBS and subjected to flow cytometry on a Becton Dickinson LSR II flow cytometer. For quantifying promoter activation from quiescence, H-Lat cells were infected as indicated and prepared for flow cytometry as described above.

### RNA extraction, cDNA synthesis, and RT-PCR

For RT-PCR analysis of *DCAF1* KD efficiency, 300,000 HeLa cells were seeded in a 6-well plate and allowed to adhere overnight. The next day, cells were PEI transfected with 300ng of pcDNA mTag-BFP2 control plasmid, pcDNA mTag *DCAF1* KD shRNA #1 plasmid, or pcDNA mTag *DCAF1* KD shRNA #2 plasmid. 72h post-transfection, cells were washed 1x with PBS, detached, and centrifuged at 500 RCF for 10 minutes. Cell pellets were resuspended in 500 uL TRizol reagent (Thermo Fisher, 15-596-018) and inubated at room temperature for 5 minutes. Next, 100 uL of chloroform was added, samples were vortexed and incubated at room temperature for 3 minutes prior to centrifugation at 12,000 x g for 15 minutes at 4 degrees. The upper phase was transferred to a fresh 1.5 mL eppendorf tube and combined with 250 uL isoporpanol for 2 hours at room temperature. Samples were centrifuged at 12,000 x g for 15 minutes at 4 degrees, pelleted were washed 1x in ice cold 70 % ethanol, and allowed to air dry. Pellets were resuspended in 20 uL RNase free H2O and immediately used for cDNA synthesis. For cDNA synthesis, 1.5 µg of RNA was incubated at 65 degrees C for 5 minutes in the presence of oligo dT primer. Then, RT buffer, dNTPs, RNase inhibitor, and reverse transcriptase were added to the reaction, incubated at 42 degrees C for 1h and then 70 degrees C for 10 minutes to inactivate reverse transcriptase. To amplify target genes, a 50 µL PCR reaction was performed following the reaction conditions described above using 0.5 uL of cDNA for 25 cycles (RT-PCR primer sequences listed below).

## Acknowledgements

This work was supported by NIAID R00 award AI147811 (to DJS), startup funds from Stony Brook University, and startup funds from the University of Texas Health Science Center at San Antonio.

## Author Contributions

Conceptualization: DJS

Funding Acquisition: DJS

Formal Analysis: DJS, NS, EL, HW, JJ, DE

Investigation: NS, EL, HW, JJ, DE

Methodology: NS, EL, HW, JJ, DE

Project Administration: DJS

Writing—original draft: DJS, NS

Writing—review & editing: DJS, NS, EL, HW, JJ, DE

---

**Figure S1. Vpr-induced global epigenetic remodeling in immortalized and primary MDMs.**

(**A**) Representative fluorescence microscopy images of histone marks in THP1 cells infected with indicated viruses.

(**B**) Representative fluorescence microscopy images of histone marks in primary MDM cells infected with indicated viruses.

**Figure S2. Vpr-induced global epigenetic remodeling in HeLa cells.**

(**A**) Representative fluorescence microscopy images and quantification of histone marks in HeLa cells infected with indicated viruses (n = 50).

(**B**) Flow cytometric analysis of DDR activation and histone marks in HeLa cells infected with indicated viruses.

(**C**) Immunoblot analysis of histone marks in HeLa cells infected with indicated viruses.

(**D** and **E**) Immunoblot analysis (D) and representative fluorescence microscopy images and quantification (E) of DDR activation in HeLa cells infected with indicated viruses.

**Figure S3. Vpr mutant-induced DDR and histone modifications**

(**A**) Quantification of DDR activation following infection of THP1 cells with Vpr_WT_ or mutant viruses (n = 50).

(**B**) Quantification of histone marks in HeLa cells infected with indicated viruses (n = 50).

(**C**) Representative fluorescence microscopy images and RT-PCR analysis of DCAF1 expression in HeLa cells transfected with DCAF1 shRNAs.

**Figure S4. Vpr modulation of HIV promoter activity**

(**A**) Flow cytometric analysis of HIV LTR activity in HeLa (top) and THP1 (bottom) cells infected with indicated viruses in the presence or absence of DDR inhibitors.

(**B**) Quantification of HIV LTR activity in primary MDMs infected with Vpr_WT_ virus in the presence or absence of DDR inhibitors.

(**C**) Quantification of histone marks in uninfected HeLa cells in the presence or absence of HDAC inhibitors (n = 50).

(**D**) Diagram displaying the establishment of the HeLa latency model H-Lats (left) and representative live cell fluorescence microscopy images (right) of H-Lat GFP expression when infected with indicated viruses.

(**E**) Flow cytometric analysis of H-Lat GFP expression when infected with indicated viruses.

